# Partial RdRp sequences offer a robust method for Coronavirus subgenus classification

**DOI:** 10.1101/2020.03.02.974311

**Authors:** David A Wilkinson, Lea Joffrin, Camille Lebarbenchon, Patrick Mavingui

## Abstract

The recent reclassification of the *Riboviria*, and the introduction of multiple new taxonomic categories including both subfamilies and subgenera for coronaviruses (family *Coronaviridae*, subfamily *Orthocoronavirinae*) represents a major shift in how official classifications are used to designate specific viral lineages. While the newly defined subgenera provide much-needed standardisation for commonly cited viruses of public health importance, no method has been proposed for the assignment of subgenus based on partial sequence data, or for sequences that are divergent from the designated holotype reference genomes. Here, we describe the genetic variation of a partial region of the coronavirus RNA-dependent RNA polymerase (RdRp), which is one of the most used partial sequence loci for both detection and classification of coronaviruses in molecular epidemiology. We infer Bayesian phylogenies from more than 7000 publicly available coronavirus sequences and examine clade groupings relative to all subgenus holotype sequences. Our phylogenetic analyses are largely coherent with genome-scale analyses based on designated holotype members for each subgenus. Distance measures between sequences form discrete clusters between taxa, offering logical threshold boundaries that can attribute subgenus or indicate sequences that are likely to belong to unclassified subgenera both accurately and robustly. We thus propose that partial RdRp sequence data of coronaviruses is sufficient for the attribution of subgenus-level taxonomic classifications and we supply the R package, “MyCoV”, which provides a method for attributing subgenus and assessing the reliability of the attribution.

**Importance Statement:** The analysis of polymerase chain reaction amplicons derived from biological samples is the most common modern method for detection and classification of infecting viral agents, such as Coronaviruses. Recent updates to the official standard for taxonomic classification of Coronaviruses, however, may leave researchers unsure as to whether the viral sequences they obtain by these methods can be classified into specific viral taxa due to variations in the sequences when compared to type strains. Here, we present a plausible method for defining genetic dissimilarity cut-offs that will allow researchers to state which taxon their virus belongs to and with what level of certainty. To assist in this, we also provide the R package ‘MyCoV’ which classifies user generated sequences.

## Introduction

Coronaviruses are widely studied for their impact on human and animal health (1) as well as their broad diversity and host/reservoir associations. In recent years, the emergence of Betacoronaviruses in human populations has resulted in widespread morbidity and mortality. The Severe Acute Respiratory Syndrome (SARS) Coronavirus was responsible for 8 096 cases and 774 deaths during the 2002-2003 outbreak (World Health Organisation, WHO data). Since 2013, the Middle East Respiratory Syndrome (MERS) Coronavirus has infected 2 506 and has led to 862 deaths (WHO data). At the time of writing, the SARS-CoV-2 virus (also known as nCoV-2019 and causing the disease Covid-19) epidemic is ongoing and is regarded as a public health emergency of international concern by the WHO, having resulted in more than 2,500 deaths. Thanks to molecular epidemiology studies, we know that SARS, MERS and SARS-CoV-2 had their origins in wild animal reservoir species before spilling over into humans. Indeed, numerous molecular studies have identified a wealth of Coronavirus diversity harboured by equally diverse animal hosts (1–6), and phylogenetic analysis of sequence data from these studies is helping in our understanding of many aspects of disease ecology and evolution (7–9). This includes the role of reservoir hosts in disease maintenance and transmission (3, 4, 6, 10–12), the evolutionary origins of human-infecting coronaviruses (3, 13), the importance of bats as reservoirs of novel coronaviruses (14, 15), the role of intermediate hosts in human disease emergence (3, 11, 16), and the understanding of risk that might be related to coronavirus diversity and distributions (17, 18).

The precise taxonomic classification of all organisms undergoes constant drift as new discoveries are made that inform their evolutionary histories. However, in 2018, the International Committee for the Taxonomy of Viruses (ICTV) introduced a shift in the taxonomic designations of all RNA viruses, introducing the realm *Riboviria*, grouping *“all RNA viruses that use cognate RNA-dependent RNA polymerases (RdRps) for replication”* (19). In addition to this basal classification, many new taxonomic classifications were defined, or existing taxa reclassified. This included the separation of the family Coronaviridae into two subfamilies – the amphibian-infecting Letovirinae, and Orthocoronavirinae encompassing the genera Alphacoronavirus, Betacoronavirus, Gammacoronavirus and Deltacoronavirus that are classically recognised to infect mammals and birds and have significance in human and livestock diseases. The subgenus level of classification for members of the Orthocoronavirinae was also introduced, providing specific taxa for commonly cited groups of similar viruses such as Betacoronavirus lineages β-A, β-B, β-C and β-D, which were designated Embecovirus, Sarbecovirus, Merbecovirus and Nobecovirus, respectively. The nomenclature for the designated subgenera was assigned with respect to known host species for each subgenus (eg. Rhinacoviruses for Alphacoronviruses known to be hosted by bats of the Rhinolophidae family) or based on commonly used disease terminology (eg. Merbecoviruses for Betacoronaviruses related to MERS Coronavirus). Whole genome data from holotype specimens selected to represent an exhaustive spectrum of coronavirus diversity was used to test the phylogenetic repartition and support for each of these taxa (20). Due to the limited diversity of holotype specimens classified into these subgenera, there is currently no method for attributing subgenus to isolates with divergent sequences, and no proposed method for partial sequence data. However, sequence data from the RdRp region of the polymerase gene is one of the most commonly used tools for the purposes of Coronavirus detection, identification and classification in molecular epidemiology.

Here, we examine the phylogenetic relationship from all identifiable public partial RdRp sequences of coronaviruses using Bayesian inference in BEAST2 and examine the clade-associations of all defined subgenus holotypes. We use this analysis to explore the range of logical similarity thresholds for the designation of subgenus-level classifications to partial RdRp sequence and predict “most-likely subgenus” classifications for all reference sequences. We cross-validate a sequence-identity-based classification method against phylogenetically inferred classifications showing that alignment identity is >99% specific for the assignment of subgenus-level classifications to partial RdRp sequences. We compiled a database of our assigned classifications and developed the R package “MyCoV” for assignment of user-generated sequences to these taxa.

## Methods

### Sequence data, curation and alignment

Sequence data was obtained from the NCBI nucleotide database on the 5^th^ of July 2019, using the search term “coronavir*”. This resulted in the identification of 30,249 sequences. A preliminary set of representative partial RdRp sequences was compiled with reference to recent publications describing Coronavirus diversity across the Orthocoronavirinae subfamily (21), in order to include starting reference sequences from with the largest possible diversity of coronaviruses. This preliminary list was then used to identify partial RdRp sequences from retrieved NCBI records by annotating regions that had at least 70 % identity to any reference sequence in the Geneious software package (version 9.4.1). Annotated regions and 200 bp of flanking sequence data were then extracted. Data containing incomplete sequences in the form of strings of N’s or significant numbers of ambiguities (>5) were removed. Open reading frames with a minimum length of 300 bp were identified and extracted from the remaining sequences. In the case where the correct reading frame was ambiguous, pairwise alignment to reference sequence data was used to determine reading frame. Remaining sequences were then aligned in-frame using MAFFT, and the resulting alignment was further curated by visual inspection. Retained sequences were then trimmed to include only the most-frequently sequenced partial region of RdRp and so that each sequence contained a minimum of 300 gap free bases. The final alignment was 387 bp in length with 7,544 individual sequences, of which 3,155 were unique. The relevant 387 bp region corresponds to nucleotide positions 15287:15673 in Merbecovirus holotype reference sequence JX869059.2.

### Genetic analyses

Phylogenies were inferred from all unique sequences using the BEAST2 software (22). Parameters were estimated for a GTR substitution model with four gamma categories and an estimated proportion of invariant sites. The Yule population model was used, and a log-normal distribution was specified for birth rate and proportion of invariant site priors. Convergence of estimated parameters was assessed in Tracer v1.7.1 (23). Three independent MCMC chains were run until effective sample sizes were above 200 for all estimated parameters after removing the burn-in. Analyses were run until convergence criteria could be fulfilled whilst providing equal chain lengths after burn-in for all three repeats, meaning that the number of trees in the posterior distributions was the same for each independent repeat.

Genetic distance measures were calculated using the ‘ape’ package (24) in RStudio as the proportion of variant sites in pairwise comparisons after removing regions containing gaps in either compared sequence.

### Taxonomic classification

Sequences originating from known references were used to identify common ancestral nodes for the *Orthocoronaviridae* genera within each phylogenetic tree. Genus-level subtrees were then extracted and treated independently for subgenus-level analyses.

Sequences originating from defining subgenus holotype samples were identified in the genus-level topologies. Clustering thresholds were defined as the highest node positions at which clusters of leaves could be defined without combining holotype specimens from different subgenera into the same clade. Clusters defined at these thresholds that contained no holotype specimens were designated as “Unclassified”. Clustering thresholds were calculated, and subgenera were assigned to all sequences across a random subsample of 453 trees, 151 from each independent repeat of the phylogenetic analyses. The proportion of trees in which each sequence was assigned to a given subgenus was used as the “posterior probability” of that sequence belonging to that subgenus. Sequences with lower than 90% majority posterior probabilities were designated as “Unclassified”. Potential positioning of new subgenus level clades (as indicated by “GroupX” in Figures 2 and 3) was inferred using the maximum clade-credibility consensus tree from all BEAST analyses, identifying monophyletic clades where all descendants were not classified into defined subgenera.

**Figure 1:**
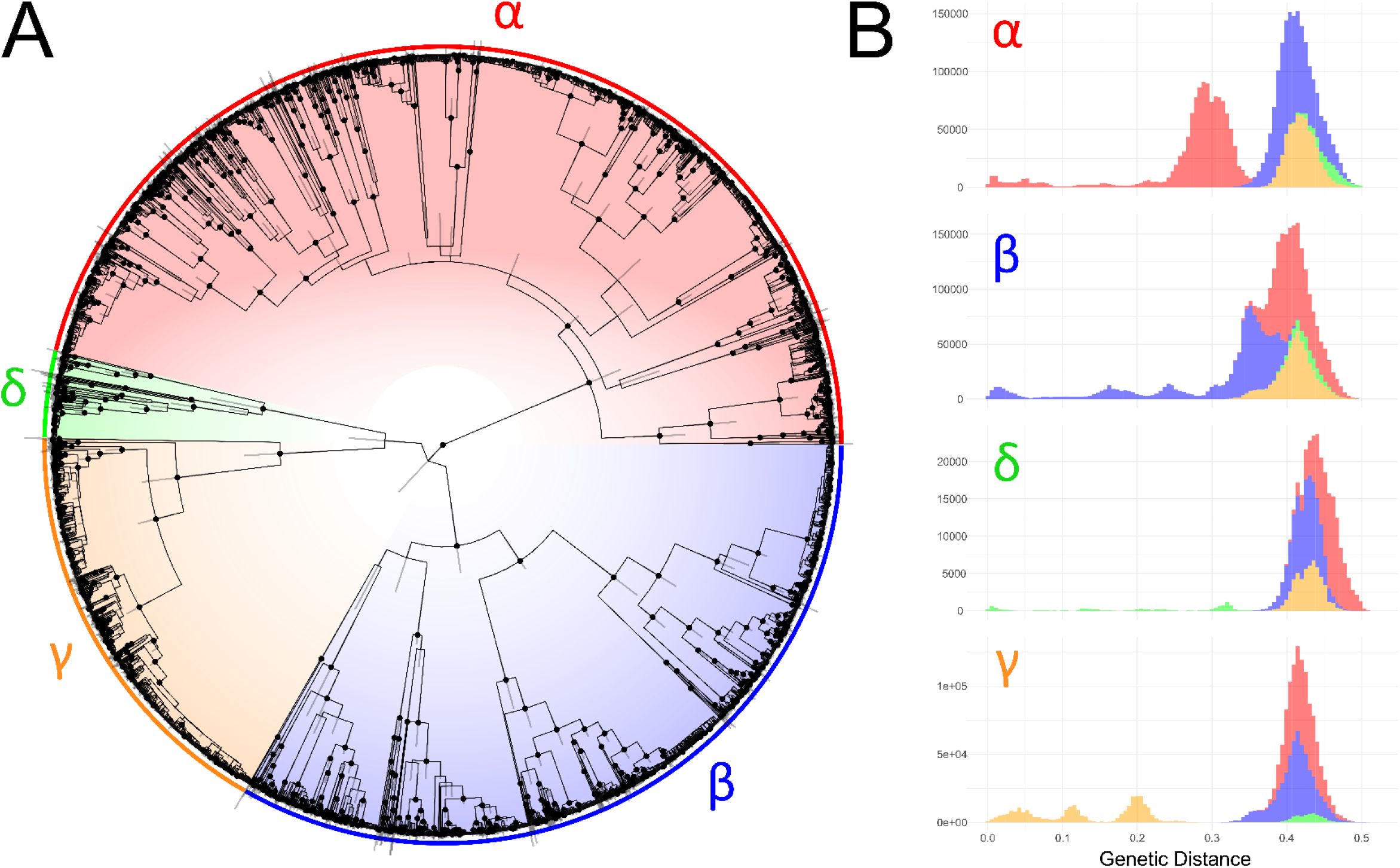
Comparative analysis of 3155 partial RdRp sequences belonging to members of the Orthocoronavirinae. A) Consensus phylogeny from three independent BEAST analyses. Nodes with posterior support greater than 90 % are highlighted with dots and bars display the 95% HPD of the heights of each node. Colours indicate genus-level classification for sequences, clades and pairwise comparisons throughout. B) Histograms of genetic distances, measured as the proportion of variant sites, between sequences belonging to each genus and grouped by the genus of the queried sequence.

**Figure 2:**
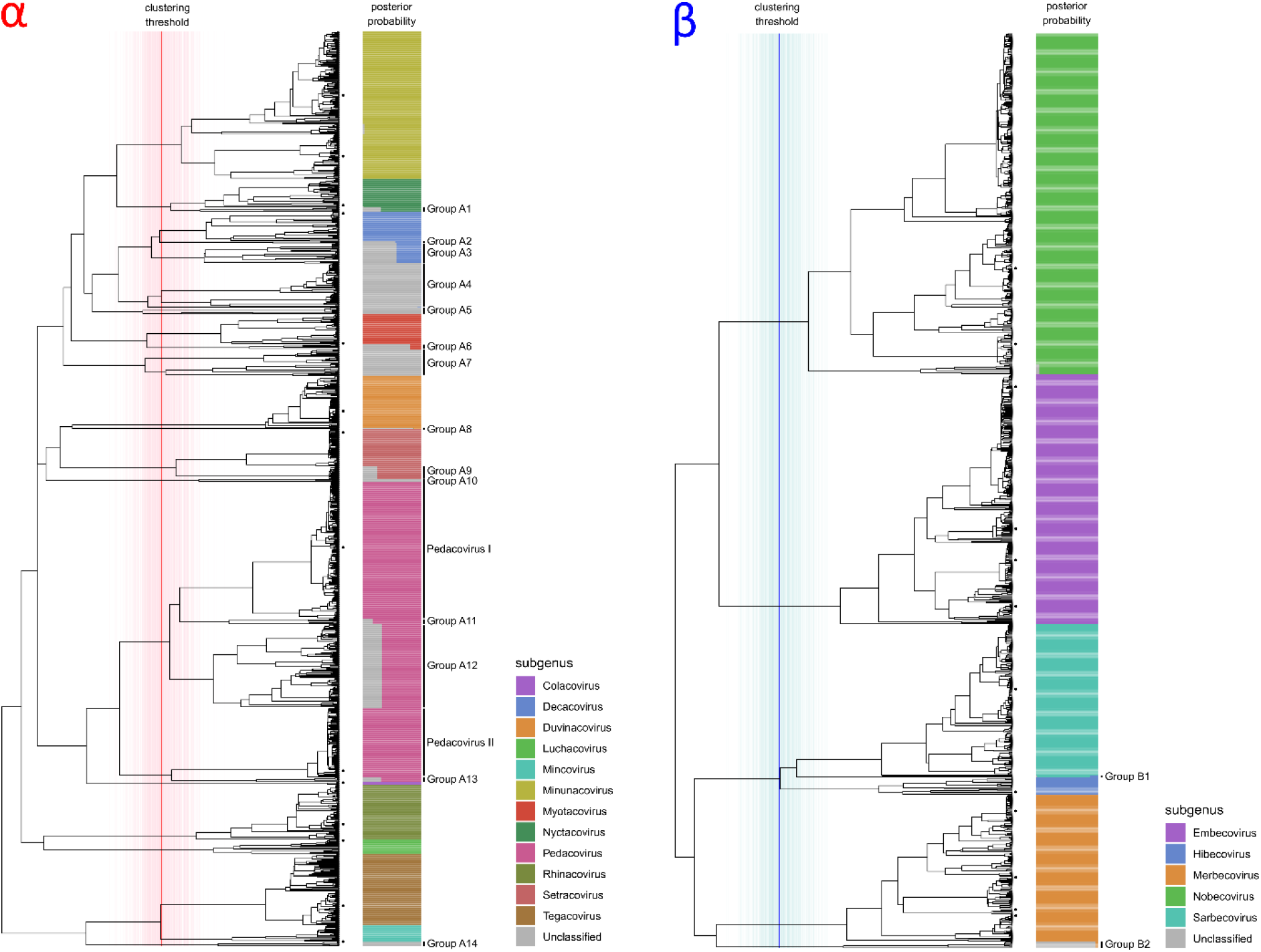
Phylogenetic subgenus classifications for partial RdRp sequences of Alphacoronaviruses (LEFT) and Betacoronaviruses (RIGHT). Depicted trees are subtrees of consensus phylogeny presented in Figure 1. Dots on leaf tips indicate sequences belonging to holotype reference sequences for each subgenus. Coloured bars show the proportion of trees from the Bayesian analysis where the corresponding leaf was assigned to each subgenus, which is indicated by colour according to the legend. Vertical lines show the distribution of cluster-defining height thresholds that were identified to assign subgenus classifications, with the median of all clustering thresholds displayed in bold, lines are coloured by genus as in Figure 1. Monophyletic groups where all members have posterior probabilities of being assigned to a known subgenus of lower than 90% are highlighted and assigned sequential IDs.

**Figure 3:**
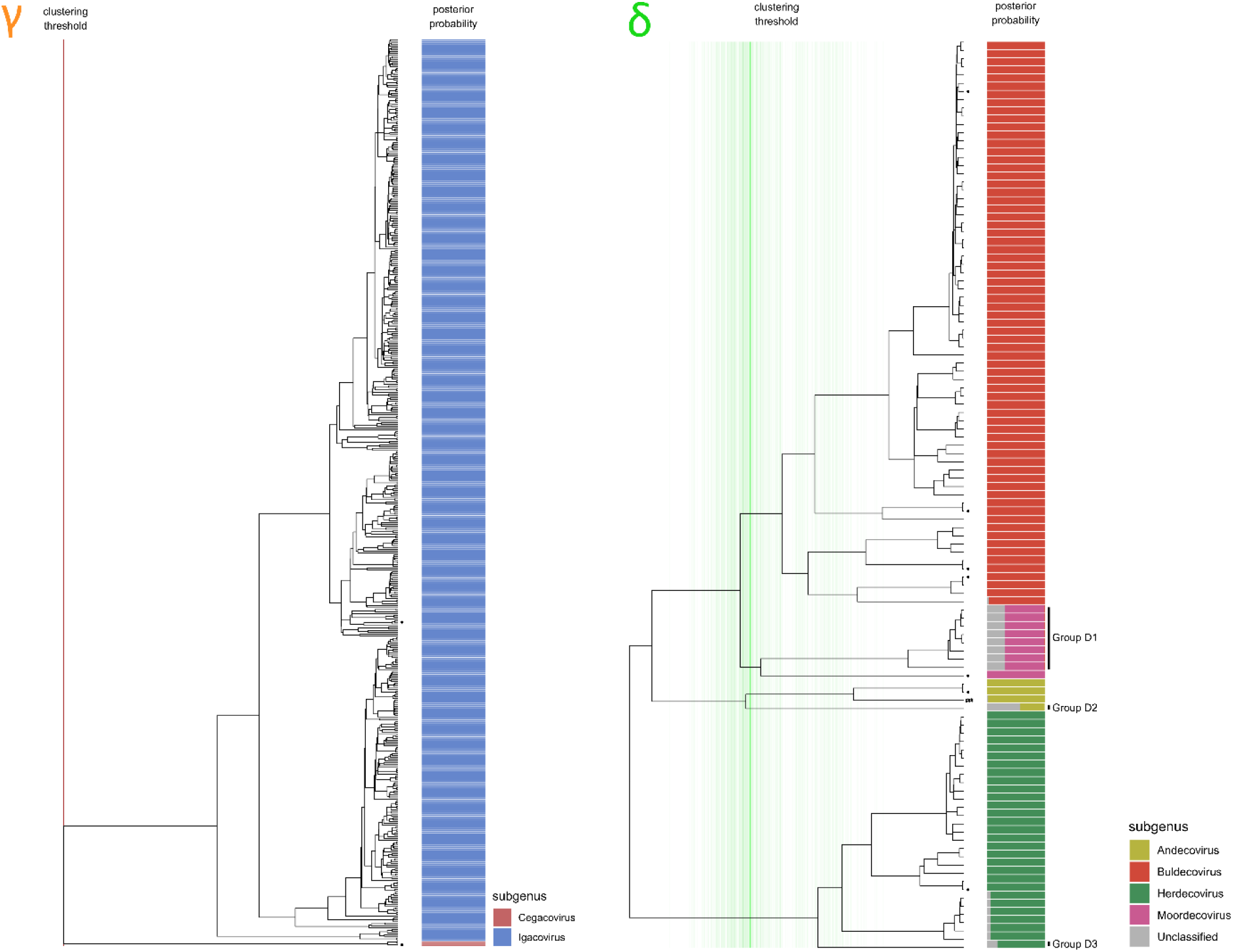
Phylogenetic subgenus classifications for partial RdRp sequences of Gammacoronaviruses (LEFT) and Deltacoronaviruses (RIGHT). Depicted trees are subtrees of consensus phylogeny presented in Figure 1. Dots on leaf tips indicate sequences belonging to holotype reference sequences for each subgenus. Coloured bars show the proportion of trees from the Bayesian analysis where the corresponding leaf was assigned to each subgenus, which is indicated by colour according to the legend. Vertical lines show the distribution of cluster-defining height thresholds that were identified to assign subgenus classifications, with the median of all clustering thresholds displayed in bold, lines are coloured by genus as in Figure 1. In the case of gamma coronaviruses, both subgenera are consistently separated at the root of the tree, thus all cluster defining heights equate to the root. Monophyletic groups where all members have posterior probabilities of being assigned to a known subgenus of lower than 90% are highlighted and assigned sequential IDs.

### Cross-validation

The assignment of sequences to the relevant subgenus using best hit and pairwise identity data from blastn (25) was tested by iteratively removing each sequence from the test database and re-assigning its classification. Sequences that could not be re-assigned to the same subgenus by this method were re-classified as “atypical” members of their respective subgenera.

## R package for assignment of user-generated sequences

The purpose of the R package MyCoV is to allow users to classify Coronavirus sequence data that includes the relevant portion of the RdRp gene to the taxonomic level of subgenus, and to assess to what extent the classification is optimal based on the criteria presented herein.

In order to achieve this, the 3155 unique partial sequences from the phylogenetic analyses were used to establish a reference BLAST database. Metadata pertaining to host organism, country of origin and date of collection were mined from NCBI and standardised by taxonomic grouping of the host and geographical region of origin to generate corresponding metadata for all 7544 NCBI reference sequences from which the unique sequence list was established.

MyCoV was written as a basic wrapper script for blastn, which queries sequences of interest against the established database, and summarises subgenus classification, subgenus posterior support of the most similar sequence in the phylogenetic analysis, pairwise distances to the most similar sequence in the database and their metadata using R packages “ggplot2”, “formattable” (available at https://github.com/renkun-ken/formattable) and “ggtree” (26).

As the MyCoV database was established prior to the recent emergence of the SARS-CoV-2, we used genomic sequence data from this virus as a test case for the utility of the MyCoV package. Outputs from this analysis are shown in Figure 5.

MyCoV is available at https://github.com/dw974/MyCoV.

## Results

Our three independent, randomly-seeded phylogenetic analyses converged on similar estimates for all parameters in BEAST2. The resulting predictions of tree topology had well supported major nodes with narrow posterior distributions around most node heights (Figure 1a). The four known genera associated with these sequences fell into four well-supported clades, divided close to the root of the tree. Genetic distance measures between all members of the four genera had logical thresholds for the distinction between genera except in the case of some betacoronaviruses, which had major clade divisions close to the root of the tree (Figure 1b) and therefore had genetic distances between members of the same genus that overlapped with distances between members of the Alpha-and Betacoronaviruses. In practice, this is likely to mean that identity-based phylogenetic topologies based on this partial region of RdRp may incorrectly infer paraphyly between members of the Alpha-and Betacoronaviruses.

At the subgenus level, separation of the inferred tree topologies into monophyletic clades based on the positions of reference holotype sequences produced logical and well-supported groupings that covered the majority of coronavirus diversity explored to date by RdRp sequencing (Figures 2 and 3). In total, 88 % of unique sequences fell into clade groups containing subgenus holotypes with subgenus-assignment posterior probabilities of greater than 90 %. The remaining 12 % of unique sequences fell into 19 separate monophyletic groups, of which 14 were Alphacoronaviruses (Figure 2), two were Betacoronaviruses (Figure 2) and three were Deltacoronaviruses (Figure 3). When host and geographical origins of isolates falling within unclassified clades was examined, the majority were associated with regional radiations for which little or no genomic or phenotypic data are available. For example, unclassified Deltacoronaviruses were all from bird species in Oceania, and many unclassified Alpha-and Betacoronaviruses originated in bat species that are exclusively found in Central and South America (Supplementary Figure 1).

The Pedacoviruses, for which multiple genome holotypes were supplied for the description of subgenus, were split into multiple clades by the imposition of a common height threshold for cluster definition using the presented methodology. The two holotype-containing clades corresponded to a single group of Porcine epidemic diarrhoea virus (PEDV) – related viruses distributed globally but entirely from pigs (Pedacovirus I in Figure 2) and a monophyletic group of viral sequences obtained from Asian *Scotophilus* bats. The monophyletic group that contained both Pedacovirus holotypes also enclosed other major viral clades (Clades A11, A12 and A13 in Figure 2), which were mainly associated with other bat species of the *Vespertillionidae* family (Supplementary Figure 1).

Cross-validation of sub-genus assignments by best-hit using blastn was successful in more than 99.9 % of cases, with a handful of lone sequences that branched at basal positions of each phylogenetic clade group being assigned to different subgenera.

For sequence members of each genus, genetic distance measurements between and within sequences attributed to each subgenus displayed logical and discrete threshold boundaries for the distinction of individual subgenus members. The one exception, again, was members of the pedacovirus subgenus which displayed overlapping within-taxon distances with between-taxon distances for other Alphacoronavirus subgenera (Figure 4). The distinct pedacovirus clades displayed in Figure 2 were thus treated as separate subgenera for distance threshold calculation. Optimal thresholds were identified as the midpoint of a fitted binomial probability distribution for intra-and inter-subgenus pairwise distances. The optimal identity thresholds for distinguishing same vs. different subgenera were as follows; i) 77.6 % identity, resulting in 99.7 % precision and 95.3 % accuracy of classification for subgenera of the Alphacoronaviruses. ii) 71.7 % identity, resulting in 99.9 % precision and 99.6 % accuracy of classification for subgenera of the Betacoronaviruses. iii) 74.9 % identity, resulting in 98.8 % precision and 99.2 % accuracy of classification for subgenera of the Deltacoronaviruses, and iv) 69.9 % identity, resulting in 100 % precision and 100 % accuracy of classification for subgenera of the Gammacoronaviruses.

Our R package, MyCoV, successfully identified SARS-CoV-2 as a member of the sarbecovirus subgenus, with the closest match being to reference sequence KP876545.11 (Rhinolophus bat coronavirus BtCov/3990), which showed 92.5 % pairwise identity to SARS-CoV-2 in the RdRp region. This sequence had been assigned 100 % posterior support for being attributed to the sarbecovirus subgenus (Figure 5a). Distributions of pairwise identities within members of the subgenera of the Betacoronaviruses fell between 71 % and 100 %, whereas pairwise distances between Betacoronavirus subgenera were less than or equal to 71%. Thus, the output of MyCoV allows us to state with certainty that SARS-CoV-2 belongs to this subgenus (Figure 5b). Positioning of the closest match in the phylogenetic tree shows that the SARS-CoV-2 forms a distinct lineage from SARS coronavirus, and that its closest match belonged to a Rhinolophus bat from China (Figure 5c). Interestingly, this sequence came from an abandoned mine in 2013, suggesting that SARS-CoV-2 predecessors circulated in bat communities for a number of years prior to the 2019 emergence in human populations. The provided visualisation of host and geographical origins for these partial reference sequences allows for a rapid assessment of the distribution of similar viruses, for example, it highlights the fact that SARS-related and SARS-CoV-2-related viruses have also been identified in bats in Africa (specifically Rhinolophus bats in Kenya), and that they are not just restricted to Asian bat hosts.

**Figure 4:**
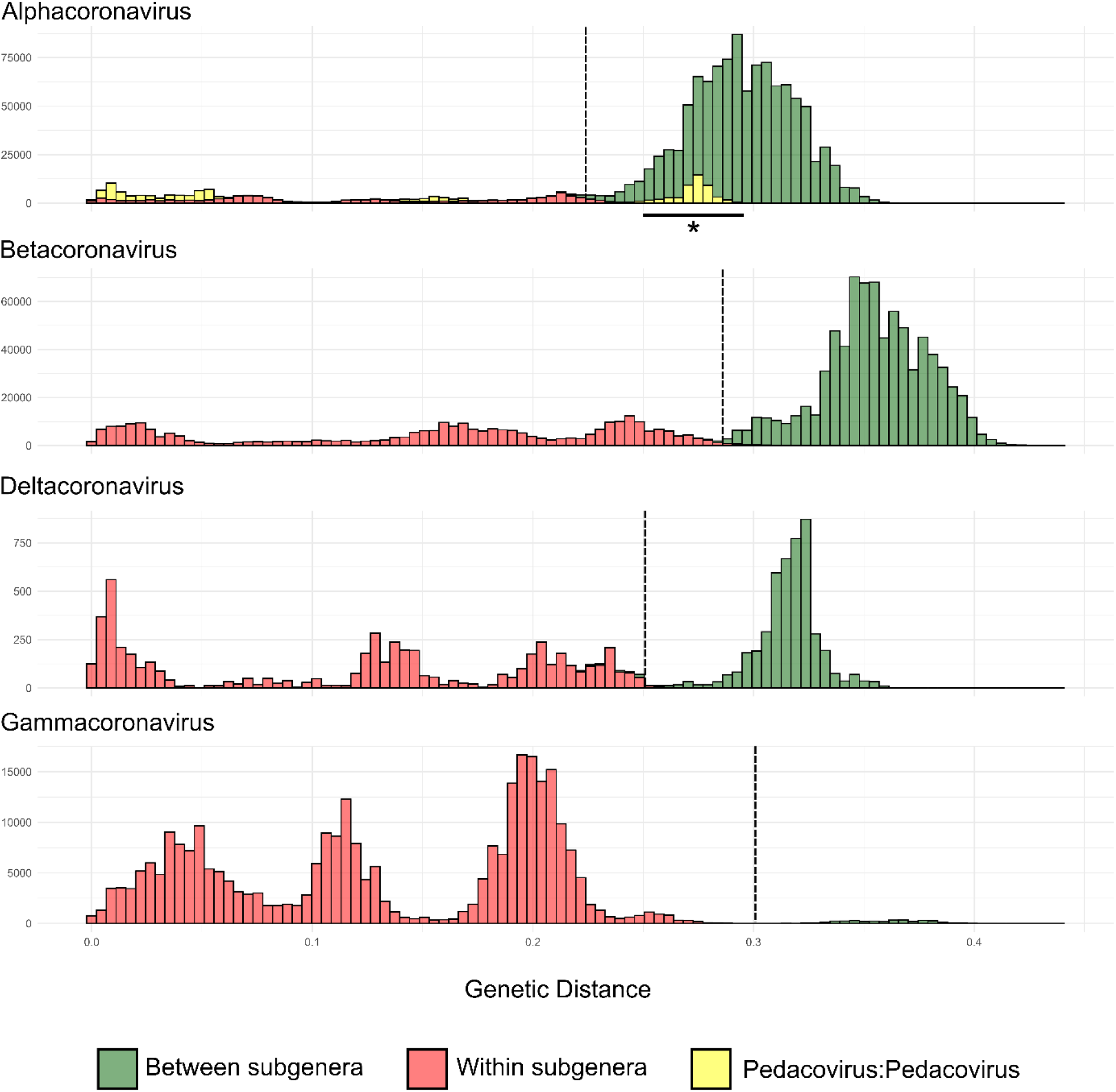
Histograms of genetic distances for intra-and inter-subgenus comparisons. Vertical dashed lines represent the optimal genetic distance cut-offs for the subgenus threshold, calculated as the midpoint of the fitted binomial probability distribution.

**Figure 5:**
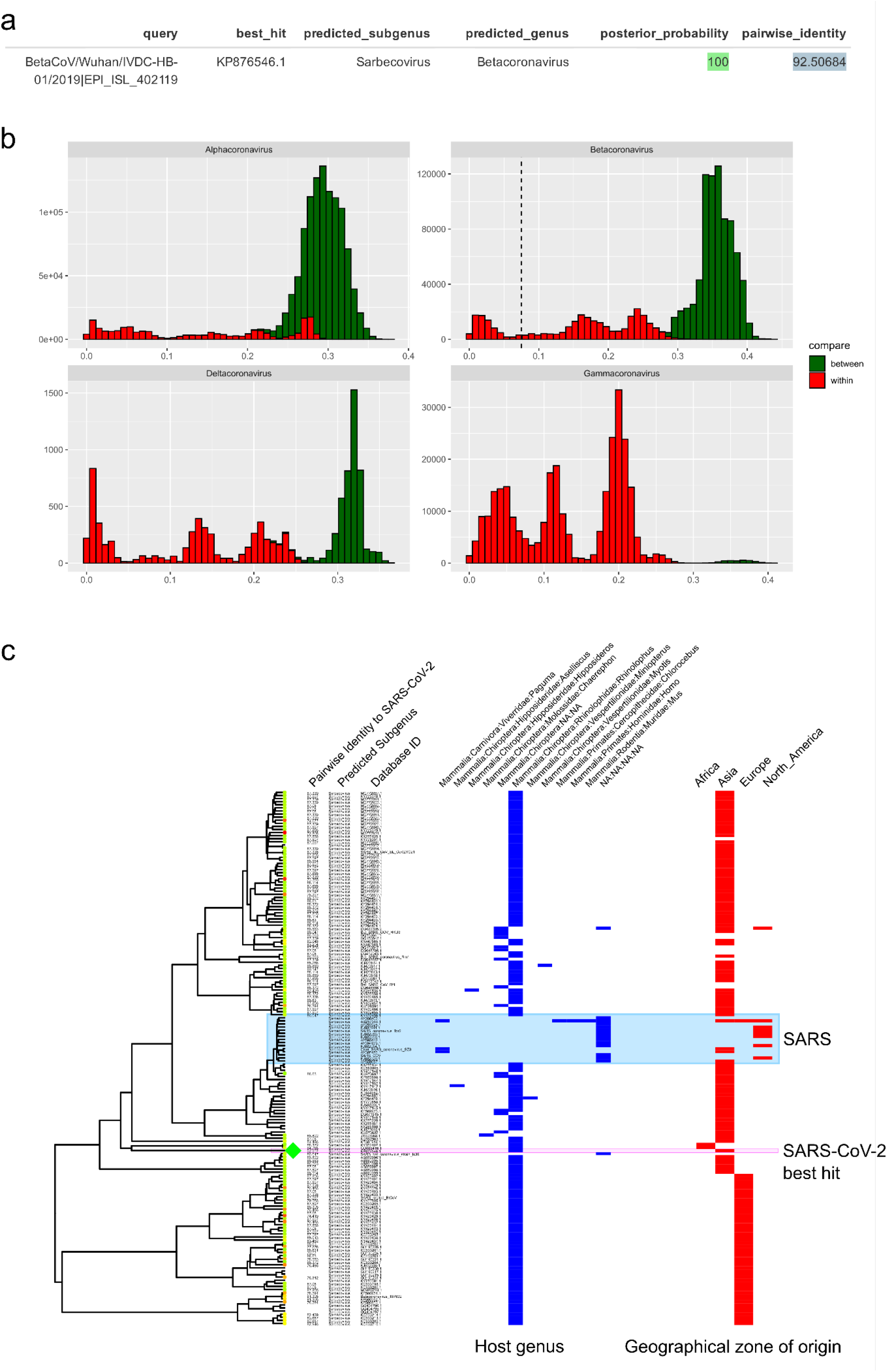
MyCoV output plots for the analysis of the 2019nCoV. A) Tabular output of best blastn hit of the query sequence to the reference database. The predicted subgenus and genus of the best-matching hit are displayed, as well as the posterior support for the assignment to the predicted subgenus (see methods). Pairwise identity between the two sequences is shown and is calculated relative to the maximum possible alignment length against the reference sequences (387 bp). B) For each queried sequence, pairwise identity values are mapped to all observations from pairwise comparisons between sequences in the database. The vertical dashed line represents the pairwise dissimilarity of the queried sequence. C) Phylogenetic positioning and metadata from the analysis of the reference sequences are displayed. Reference sequences with blast-hits matching the queried sequence are highlighted on the leaves, and tips are coloured from red to green with increasing pairwise identity. The hit with the best score is highlighted by a large green diamond on the tip. Pairwise identity scores are displayed for all leaves, as well as predicted subgenus. Host genus associations (blue) and geographical region of origin (red) from available metadata are indicated by binary heatmaps. Note that multiple metadata observations are possible for each leaf, as leaves are displayed for unique sequences only. The ID next to each leaf is that of the representative sequence for that leaf, and other IDs are left off for clarity.

## Discussion

The recent reclassification of the *Riboviria* is a logical progression in viral taxonomy, as the unique mechanism of replication of all negative-sense, single-stranded, RNA viruses results in the conservation of many viral characteristics, including relative sequence conservation of regions of the cognate RNA-dependent RNA polymerase. Consequently, such genomic loci lend themselves to the design of primers for virus detection in diagnostics and molecular epidemiology, and to the phylogenetic inference of evolutionary histories. Furthermore, establishing the classification level of subgenus has provided a useful tool for researchers, attributing standardised terminology for many commonly referenced viral lineages that, in general, demonstrate a level of specificity in their host-associations and epidemiological characteristics (Supplementary Figures S1).

Our analyses have shown that the phylogenetic interpretation of short sequences of the RdRp locus of members of the *Orthocoronaviridae* is largely coherent with genome-scale analyses based on designated holotype members for each subgenus. The vast majority of known RdRp sequences (88 %) can be classified into the defined subgenera, and their classification cross-validated based on simple distance thresholds established from a 387 bp fragment of RdRp. Globally, these distance measures form discrete clusters between taxa, offering logical threshold boundaries that can attribute subgenus or indicate sequences that are likely to belong to unclassified subgenera both accurately and robustly without the need for complex phylogenetic inference. The provided R package, “MyCoV”, provides a method for achieving this and for the assessment of the reliability of the attribution.

An alternative strategy for coronavirus classification from partial sequence data may be using the spike protein-encoding S-gene, which is another commonly sequenced region of coronavirus genomes. However, the use of this region is more common in epidemic outbreak scenarios and thus there are many S gene sequences in public databases that are either identical or extremely closely related. Performing comparative sequence searches by querying the NCBI nucleotide database with the two search terms “((coronavir* spike) OR (coronavir* S gene)) AND “viruses”[porgn:_txid10239]” and “((coronavir* RdRp) OR (coronavir* polymerase)) AND “viruses”[porgn:_txid10239]” shows that there are approximately three times more sequences from the S gene, but that these sequences originate from approximately three times fewer viral taxa. We therefore favour the use of the RdRp region as it provides a more exhaustive representation of known coronavirus diversity.

Of course, this form of interpretation is subject to the same caveats as any other that is based on partial sequence data from a short, single genomic locus; Indeed, the effects of potential recombination events cannot be captured, and some uncertainties will exist in the presented phylogenetic trajectories that may be resolvable by the addition of longer sequence data. For these reasons, we do not suggest the definition of new subgenera for unclassified clade groups presented in Figures 2 and 3. The limits of the phylogenetic resolving power of this partial region of RdRp are most clear for members of the Alphacoronavirus genus, where there is an elevated level of mid-distance genetic diversity and a large number of unclassified genetic clade groups associated with regional, likely host-specific radiations. And thus, precise taxonomic delineation of emerging Alphacoronaviruses will require more information than is offered by this RdRp locus. Conversely, the clear genetic distinction and corresponding epidemiological associations that exist between clade groups of the Pedacoviruses does raise the question as to whether the definition of this subgenus should be revisited.

---

**Supplementary Figure S1:**
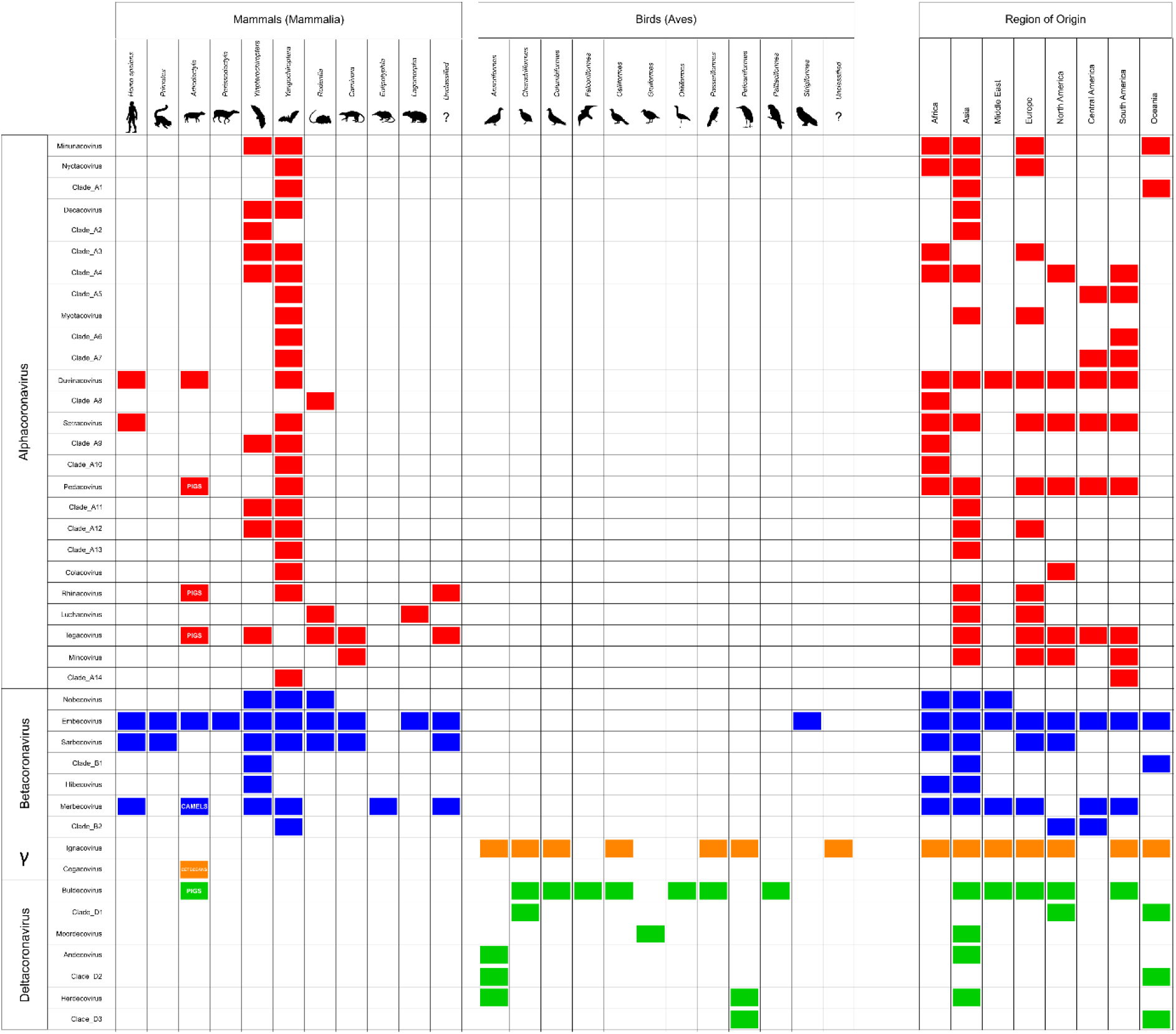
Metadata (host and geographical origin) associations of Coronaviruses belonging to different official subgenera, and other unclassified major clade groups as depicted in main figures 2 and 3. Blocks of colour represent the existence of at least one record of the indicated association, and are coloured by viral genus as in main figure 1.

